# Differential chromatin accessibility landscape of gain-of-function mutant p53 tumours

**DOI:** 10.1101/2020.11.24.395913

**Authors:** Bhavya Dhaka, Radhakrishnan Sabarinathan

**Affiliations:** National Centre for Biological Sciences, Tata Institute of Fundamental Research, Bengaluru 560065, India

## Abstract

Mutations in TP53 not only affect its tumour suppressor activity but also exerts oncogenic gain-of-function activity. While the genome-wide mutant p53 binding sites have been identified in cancer cell lines, the chromatin accessibility landscape driven by mutant p53 in primary tumours is unknown. Here, we leveraged the chromatin accessibility data of primary tumours from TCGA to identify differentially accessible regions in mutant p53 tumours compared to wild p53 tumours, especially in breast and colon cancers. We found 1587 lost and 984 gained accessible regions in breast, and 1143 lost and 640 gained regions in colon. However, less than half of those regions in both cancer types contain sequence motifs for wild-type or mutant p53 binding. Whereas, the remaining showed enrichment for master transcriptional regulators, such as FOX-Family TFs and NF-kB in lost and SMAD and KLF TFs in gained regions of breast. In colon, ATF3 and FOS/JUN TFs were enriched in lost, and CDX family TFs and HNF4A in gained regions. By integrating the gene expression data, we identified known and novel target genes regulated by the mutant p53. Together, these results suggest the tissue- and tumour-type specific role of mutant p53 in regulating chromatin structure and gene expression.

## Introduction

The p53 protein, encoded by the TP53 gene, is a transcription factor that acts as a tumour suppressor. It is a multi-domain protein that consists of an N-terminal transactivation domain, a central DNA-binding domain (DBD), and an oligomerization domain that are required for the tetramer formation of p53, DNA binding and transactivation of target gene expression (1, 2). The activity of p53 is induced by various stress signals like DNA damage, hypoxia, oxidative stress and nutrient deficiency (3). Under these conditions, the p53 binds to a specific DNA sequence (referred to as p53 response element) and regulates the target gene expression to prevent genomic instability, inhibit cell growth or trigger apoptosis (4). In most cancers, the function of p53 is affected due to genetic aberrations (mutations, copy-number loss or/and epigenetic alterations) in TP53 or its regulators (such as MDM2) (5). The mutations targeting different domains of p53 can exert distinct effects on the p53 function such as complete or partial loss of tumour suppressor activity, dominant-negative effects, and oncogenic gain-of-function (GOF) properties (6).

Missense mutations account for >70% of all the somatic alterations observed in TP53 in human cancers, most of these are clustered as hotspots in the DBD (6, 7). DBD mutations can be classified into two main groups: *DNA-contact mutations*, which affect the amino acid residues involved in the DNA binding and *conformational mutations,* which alter the folding and structure of p53 (8). In both cases, the mutations can directly or indirectly affect the DNA-binding ability of p53. Besides, in the early stages of tumorigenesis, some of these DBD mutations also show dominant-negative effects by inactivating the function of wild-type p53. However, in later stages, when the loss of wild-type TP53 allele has occurred, these DBD mutations exhibit oncogenic GOF activities that enhance tumorigenic potential. Eventually, the cancer cells become dependent on the mutant p53 for their growth and survival (6, 9).

Unlike the wild-type p53, the GOF mutant p53 does not bind to a specific DNA sequence motif (10). Several mechanistic explanations for the GOF activity of p53 mutants have been proposed, including: a) Complex formation with cellular transcription factors, where mutant p53 contributes its active transactivation domain and thereby altering the expression of their respective target genes. Examples of such TF partners include NF-Y, SREBP-2, SP1/2, ETS1/2, PML and VDR; b) Recruitment of mutant p53 by cellular transcription factors to their cognate binding sites, leading to more robust transactivation of their respective target genes; c) DNA structure-specific binding of mutant p53 in the promoter regions, resulting in transcriptional regulation of the relevant genes, and d) Direct recruitment of mutant p53 to the regulatory regions and/or chromatin changes through interaction with chromatin-modifying enzymes (2, 11, 12).

Although the potential targets of GOF mutant p53 have been discovered based on the analysis of gene expression (13, 14), the transactivation mechanism is not fully understood (especially in the above-said context). Previous ChIP-seq experiments on GOF mutant p53 helped to identify the genomic locations bound by the mutant p53 directly or through its interacting partners (15, 16). However, changes in the chromatin landscape of the target regions and nearby regulatory elements in the primary tumours remain elusive. In this study, we leveraged the chromatin accessibility data (ATAC-seq) generated in the primary tumours from The Cancer Genome Atlas (TCGA) (especially in breast and colon cancers) and identified differential chromatin accessible regions in GOF mutant p53 tumours compared to wild-type p53 tumours. Moreover, we identified transcription factor binding sites that were specifically enriched in the gained and lost peaks of chromatin accessibility in mutant p53 tumours, and their effect on the expression of the target genes. Overall, our data suggest the direct and indirect mechanisms by which GOF mutant p53 targets the chromatin and subsequent gene expression patterns in a tumour-type specific manner.

## Results

### Identification of differential chromatin accessibility in gain-of-function mutant p53 tumours

From the TCGA cohort of 404 tumour samples (across 23 cancer types) with genome-wide chromatin accessibility data (ATAC-seq) (13), we identified 33 samples harbouring gain-of-function missense mutations (or hotspot mutations) in the DNA-binding domain of p53 (Supplementary Table S1). Most of which (28/33) exhibited biallelic alterations, that is, missense mutation on one allele and copy-number loss on another allele (consistent with Knudson’s two-hit hypothesis (17)). On the other hand, we identified 64 samples as wild-type p53, which have neither protein-affecting mutations (non-synonymous or nonsense) nor copy-number alterations in TP53 (Supplementary Table S1, see Methods). To perform differential chromatin accessibility analysis between mutant and wild-type p53 tumours, we focused on two cancer subtypes, breast infiltrating ductal carcinoma and colon adenocarcinoma, which had an almost equal number of samples in both groups. The breast infiltrating ductal carcinoma consists of six mutant samples (two R273C and one each of R273P, R273H, G245C, R175H) and five wild-type p53 samples; and colon adenocarcinoma with eight mutant samples (four R273H and one each of R282W, R248Q, R175C, R175H) and six wild-type p53 samples.

We performed differential chromatin accessibility analysis at the individual cancer subtype level to avoid confounding effects from tissue- and tumour-type variabilities (see Methods). With the |log-fold change| > 1 and FDR (q-value) < 0.1, we found 1587 lost and 984 gained peaks of chromatin accessibility in breast carcinoma (out of 209,835 peaks analysed), and 1143 lost and 640 gained peaks in colon adenocarcinoma (out of 119,925 peaks) (Figure 1A, B; Supplementary Table S2). Based on the genomic annotations, we identified that the majority of gained and lost peaks were located in intronic and intergenic regions in both breast and colon adenocarcinoma (Figure 1A, B), as expected. The unique number of protein-coding genes corresponds to 661 and 558, respectively, in lost and gained peaks of breast carcinoma; and 519 and 379, respectively, in lost and gained peaks of colon adenocarcinoma. Of these, 28 genes respectively in lost and gained peaks of breast; and 19 genes in lost and 24 genes in gained peaks in colon adenocarcinoma were known cancer drivers (from COSMIC Cancer Gene Census (18)) (Figure 1A, B).

**Figure 1:**
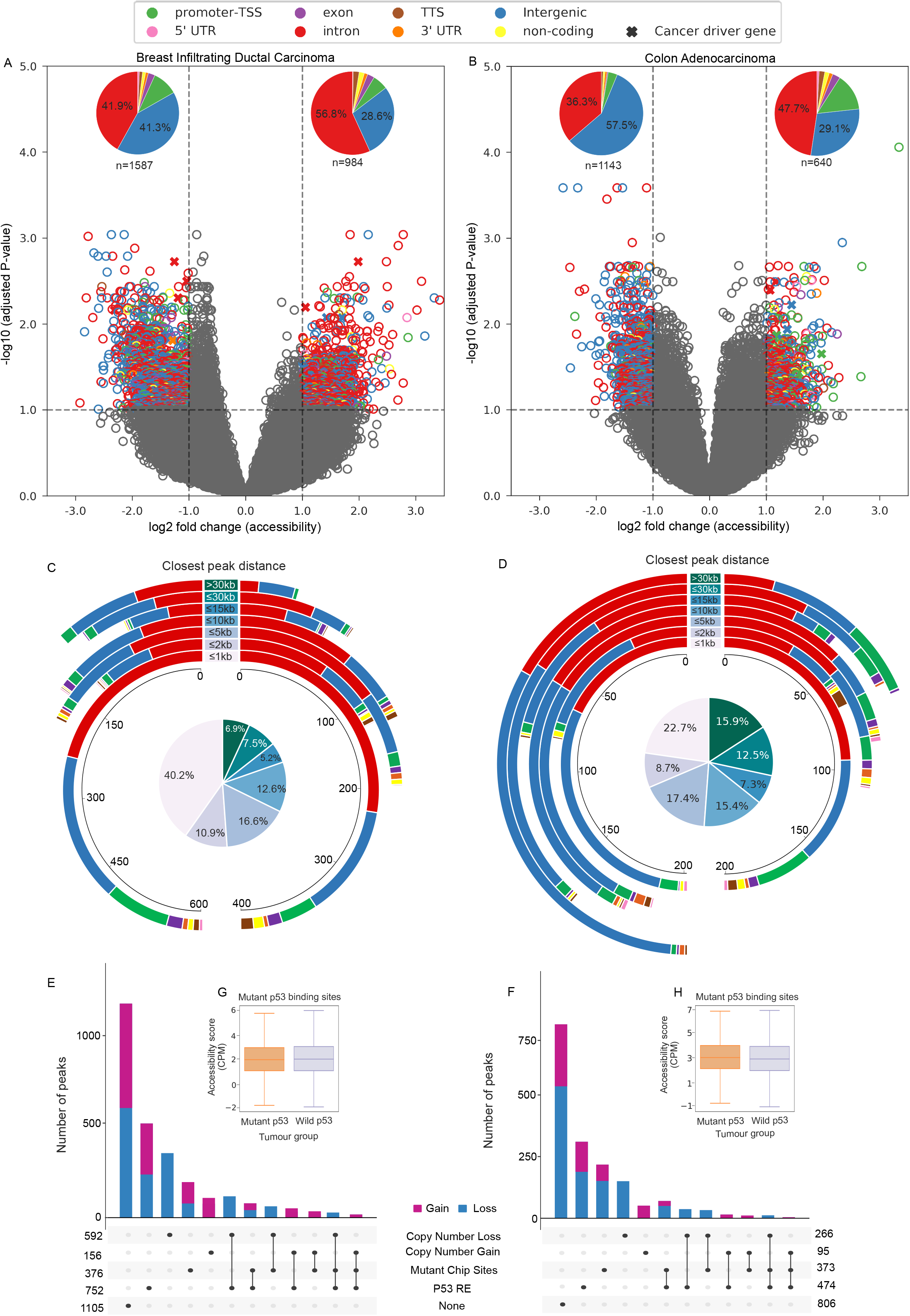
Differential chromatin accessible peaks in mutant p53 tumours. (A-B) Volcano plot shows the differential chromatin accessible peaks in the mutant p53 with respect to the wild-type p53 tumours in breast infiltrating ductal carcinoma (left) and colon adenocarcinoma (right). Each dot represents a peak region and the colour represents the genomic annotation (for significant candidates).The x-axis represents log2 fold change of accessible score in mutant versus wild-type p53 tumours, and the y-axis shows the −log10 adjusted p-value. The vertical and horizontal dashed lines indicate the threshold of FDR (q-value) < 0.1 and |log2FoldChange(Accessibility)| of 1. The symbol X indicates the peaks that overlapped cancer driver genes. The pie chart on the top represents the distribution of each genomic feature in significant lost and gained peaks, where *n* represents the total number of peaks. (C-D) The donut plot shows the distribution of closest peak distance for significant lost (left) and gained (right) peaks compared to the non-significant peaks. Each circle represents a distance bin and the stacks within that represent the peaks with specific genomic annotation (as shown in panels A and B). The pie chart inside represents the overall proportion of peaks that fall under each distance group (highlighted with the same colour code mentioned at the center of the donut plot). (E-F) Upset R plot showing distribution and overlap within different sets of annotations of the significant lost and gained peaks. (G-H) The box plot shows the median CPM (counts per million) value computed across mutant and wild-type p53 samples for all the peaks (both significant and nonsignificant) that overlapped with the mutant p53 binding sites (inferred from ChIP-seq).

Further to check if the gained and lost peaks are proximal to existing peaks (indicating the extension or reduction of already existing open chromatin region), we computed the distance of each of the gained/lost peaks against other peaks (which did not show the significant difference) in that respective cancer type. This showed that the majority of the gained and lost peaks altogether, around 40% in breast and 23% in colon adenocarcinoma, were indeed close (within 1kb distance) to the existing peaks in the respective tumours, as compared to the rest that were located at >1kb distance (Figure 1C, D). However, in colon adenocarcinoma, we observed a relatively higher number of lost peaks spread across different bins of >1kb distance when compared to gained peaks (Figure 1D). To check if this could be due to copy number alterations (CNA) in mutant or wild type p53 samples, we annotated each of the peaks having CNA gain or loss based on the CNA segmentation data from TCGA (see Methods). Overall, the lost peaks have higher overlap (23% in colon and 37% in breast) with the CNA alterations when compared to the gained peaks (15% in colon and 16% in breast), however, these were not particularly enriched in the >1kb distance bins (Supplementary Table S3).

We then sought to identify if the gained and lost peaks overlap with the known binding sites of mutant p53 (inferred from previous ChIP-Seq analysis in cancer cell lines) (15, 16) or predicted to have wild-type p53 response elements (RE) (19), both canonical and non-canonical (see Methods) (Figure 1E, F). Approximately one-fourth of the gained and lost peaks (29% in breast and 26% colon carcinomas) were predicted to have wild-type p53 RE. However, as compared to this, a relatively small proportion (15% in breast and 20% in colon carcinomas) overlapped with mutant p53 binding sites. To check this further, we compared the chromatin accessibility score for all mutant p53 binding sites in wild-type and mutant p53 tumours (Figure 1G and H). This showed that overall there was not much difference between the wild-type and mutant p53 tumours (regardless of the genomic location, Supplementary Figure S1), with few exceptions detected from our differential analysis. Taken together, these results suggest that the majority of the mutant p53 binding sites, inferred from the ChIP-seq analysis, were located in the existing open chromatin regions and that these regions did not exhibit any local chromatin structural changes when compared to wild-type p53 tumours.

In summary, we identified genomic regions that gained or lost chromatin accessibility in mutant p53 tumours compared to the wild-type p53 samples. However, only less than 50% of them have wild-type p53 response elements and/or overlap with mutant p53 binding sites.

### Enrichment of transcription factors binding motifs in the differentially accessible regions

Mutant p53 is known to interact with other TFs (such as SP1/2, ETS1/2, E2F1, ETV1, NF-Y, NF-kB, SREBP, and SMAD1/2) and histone modifiers (MLL1/2) (16) that results in altered expression of target genes. To identify which TFs are bound to the gained and lost peaks, we performed enrichment analysis of known TF binding motifs, as well as *de novo* motif discovery (see Methods). The TF motif enrichment analysis (based on motif predictions from JASPAR) exhibited that around 50-60% of gained or lost peaks overlapped with ZNF263, SP1, and SP2, and other known co-factors of p53 (as mentioned above) in both breast and colon cancers (Figure 2A-D, Supplementary Table S4), but their enrichment was not significant (q>0.1) as compared to non-significant peaks. This can be explained by the fact that some of these TFs are generally enriched in the promoter or enhancer regions. However, the TFs such as KLF5 (52%), KLF1 (39%), and SMAD2::SMAD3::SMAD4 (31%) showed significant enrichment (q<0.1) in gained peaks and FOX family of TFs (FOXA1 28%, FOXA2 28% and FOXF2 17%) in lost peaks in breast carcinomas. In contrast, colon adenocarcinoma showed a significant enrichment of TFs such as CDX2 (26.5%), CDX1 (24%), HOXA13 (22%), HNF4G (37.2%), and TCF4 (36%) in gained peaks; and FOS::JUNB (45.4%), FOS::JUND (47.5%) and JUNB (46.3%) in lost peaks.

**Figure 2:**
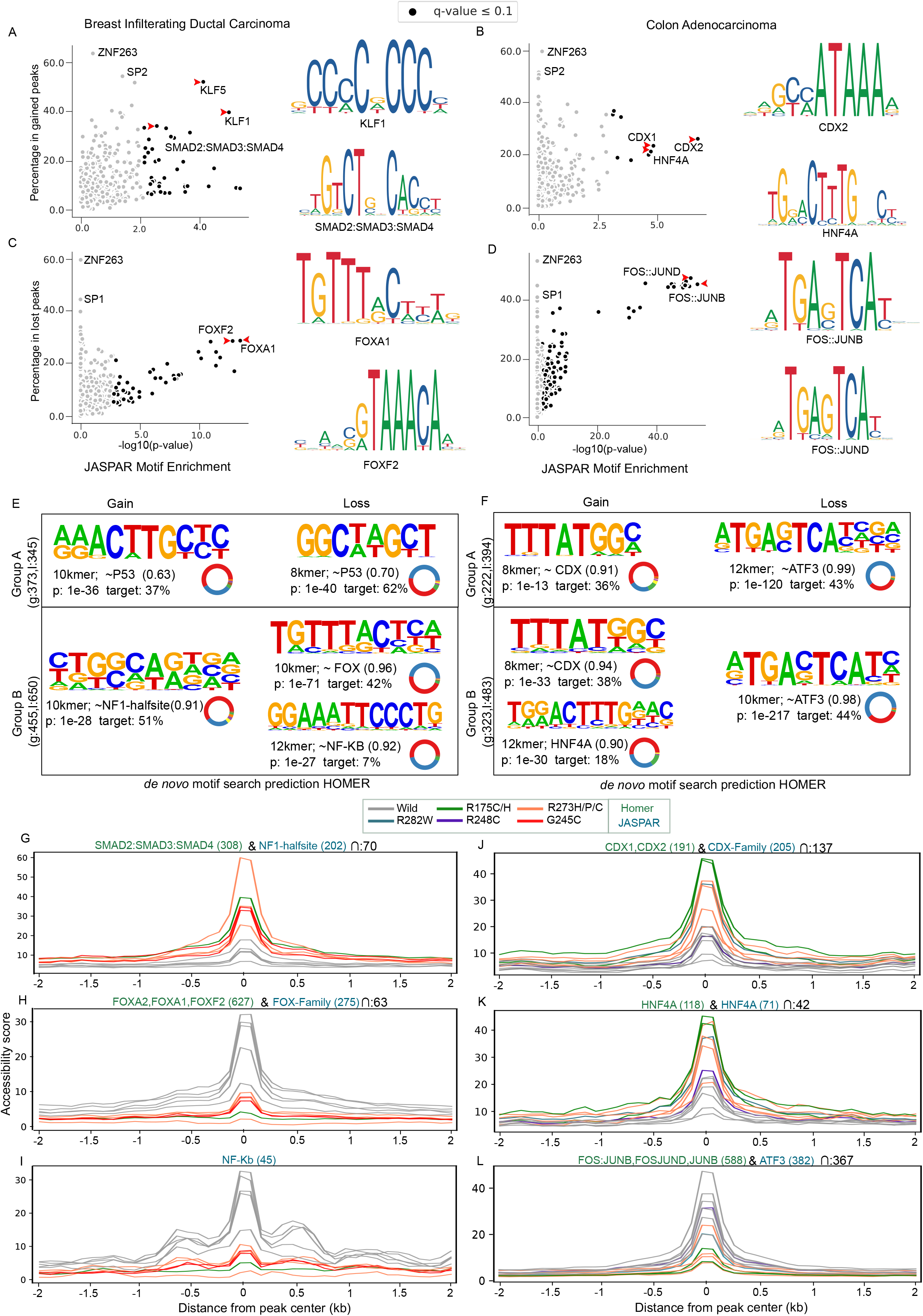
Enrichment of transcription factor motifs in differential accessible regions. (A-D) Scatter plot shows the enrichment of predicted Transcription Factor (TF) motifs (from JASPAR) in gained and lost peaks of breast and colon cancers. Each dot represents a TF, the y-axis represents the proportion of peaks overlapped with the predicted motifs of that particular TF, and the x-axis represents the significance of enrichment compared to random peaks computed using one-sided Fisher’s exact test. The TFs that were significantly enriched (q-value <=0.1) are highlighted in black. Sequence logo of a few significant TF motifs is shown beside the scatter plots. (E-F) The panels show the logo of de novo sequence motifs identified by HOMER in the group A and B peaks (see methods) and their similarity with the known TF motifs (the value close to 1 indicates high similarity). The donut plot besides it shows the distribution of genomic features in peaks that contained the particular motif. The colour code corresponds to the genomic features shown in Figure 1. (G-L) The plot shows the accessibility score at the centre of peaks (containing specific TF motif) compared to its immediate flanking regions in mutant and wild-type p53 samples. The colour of the line indicates different mutant samples and the text colour of the motif name represents whether it was found from JASPAR or HOMER de novo motif analysis.

In addition, we performed *de novo* motif discovery analysis (using HOMER) to check for the enrichment of sequence motifs in differentially accessible peaks, especially in the two distinct groups: (A) peaks with predicted p53 RE and/or overlap with mutant p53 ChIP-seq signal (discussed in the above section), and (B) the rest (see Methods, Supplementary Figure S2). The top k-mer motifs identified in each group, along with their similarity (alignment) score with known TF motifs, is shown in Figure 2E-F (and the full list of motifs identified is shown in Supplementary Table S5). In group A of breast carcinomas, we found a 10-mer motif in gained peaks (enriched in 37% of peaks) and 8-mer motif in lost peaks (62%) showing similarity with p53 motif and also partially resembling the p53 response element 5′-RRRCWWGYYY-3′ (where R = purine, Y = pyrimidine and W is either A or T), as expected. Whereas in group B, we found an 8-mer motif in gained peaks (44%) showing high similarity with NF1-half site. On the other hand, we found one 10-mer motif in lost peaks (42%) showing high similarity with FOX family and another 12-mer motif showing high similarity with NF-kB and RELA (enriched in 7% of the peaks). In colon adenocarcinomas gained peaks, we found an 8-mer motif showing high similarity with CDX family TFs in both group A (36%) and B (38%). In group B, an additional 12-mer motif showing similarity with HNF4A was found in 18% of gained peaks. Whereas in lost peaks, we found a 12-mer and 10-mer motif, respectively in group A (44%) and B (43%), showing high similarity with ATF3. Overall, the results obtained from this analysis is complementary to the above TF motif enrichment analysis and the motifs share the same genomic location (Supplementary Figure S3).

Interestingly, some of the TFs that showed high similarity with de novo motifs are the known master regulators involved in chromatin remodelling and gene regulation. For example, the FOX family of TFs, enriched in the lost peaks of breast carcinomas, is a pioneering factor which can bind to closed chromatin structure to facilitate DNA accessibility (20). To check the extent of loss of chromatin accessibility in the neighbouring regions, we compared the accessibility score at the lost peaks (with FOX family TFs motifs) with the immediate flanking regions (+/− 2kb) (Figure 2H). In wild-type p53 samples, we observed an increase in accessibility at the centre of the peaks, and also in the surrounding regions (+/− 500bp). However, in the mutant p53 samples, the accessibility score was lower at both the peak centre and flanking regions compared to the wild-type p53. This suggests that these regions are less accessible (or repressed) due to the loss of FOX family TF binding in mutant p53. Similarly, in the lost peaks enriched with NF-kB and RELA motifs, we observed a decreased accessibility at the centre of the peaks and in the flanking regions in mutant p53 compared to the wild-type (Figure 2I). In the case of colon adenocarcinoma, the CDX family TFs and HNF4A, enriched in the gained peaks, have previously been shown to bind enhancer regions and maintain chromatin accessibility (21). Consistent with this, our accessibility profile also showed a sharp increase at the peak centre and in the surrounding regions (+/− 500 bp) (Figure 2J, K). On the other hand, the TFs ATF3, FOS, JUNB and JUNC enriched in the lost peaks belong to members of the AP-1 complex. Depending on the complex formation between FOS, JUNB/C and AFT3, this can act as transcriptional activator or repressor (22). In our case, we observed a strong decrease in the accessibility for mutant p53 (Figure 2L), suggesting the repressive activity mediated by the loss of AP-1 complex at these sites.

Taken together, these results suggest that besides the enrichment of p53 response elements and mutant p53 binding sites, the master TFs (FOX, CDX and AP1) that could potentially alter the chromatin accessibility and gene regulation were enriched in the gained and lost peaks in a tissue- and tumour-specific manner.

### Enrichment of enhancer marks and G-quadruplex structure in the differentially accessible regions

To check if the gained and lost peaks overlap with the regulatory regions (such as enhancer and histone marks), we overlapped these peaks with the enhancer predictions (from GeneHancer) as well as available histone marks (such as H3K27ac associated with enhancer regions) from cancer cell-lines harbouring gain-of-function mutant p53 (see Methods). In breast carcinoma, we observed that around 50% of the peaks, in both gained or lost, overlapped with the enhancer regions. Moreover, the peaks that overlapped both enhancer prediction and H3K27ac marks in breast cancer cell lines were higher in gained peaks compared to the lost peaks (Figure 3A). Similar results of enhancer overlap were observed in colon adenocarcinoma (Figure 3B), however, the regions that overlapped with both enhancer and H3K27ac were less (except in HT-29 cell line). Interestingly, the peaks that overlapped H3K27ac (including those that overlapped both H3K27ac and enhancers) showed higher accessibility compared to peaks that overlapped only the enhancers or none (Figure 3C, D).

**Figure 3:**
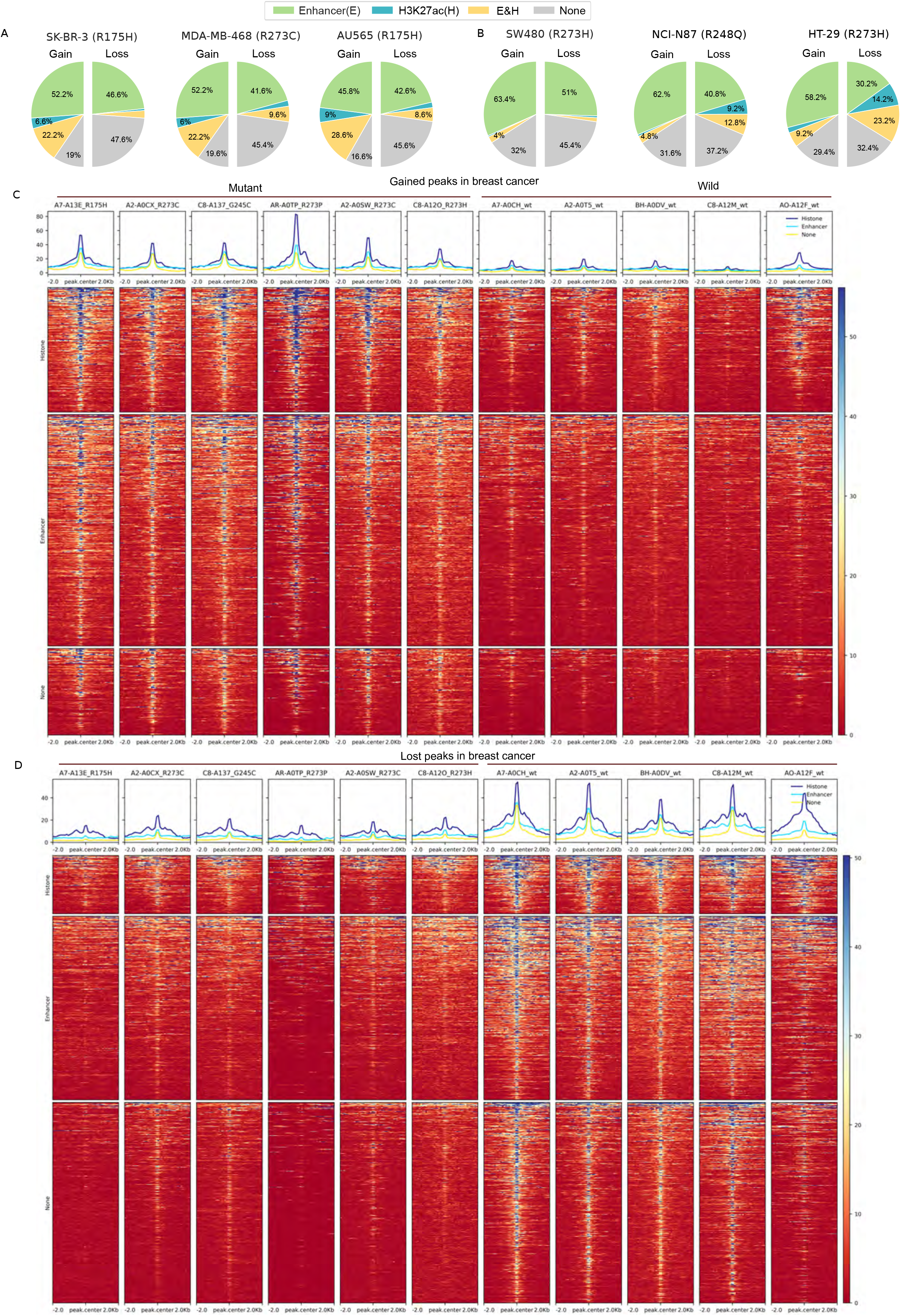
Enhancer and histone marks overlap with the differential accessible peaks. (A-B) The pie-chart shows the proportion of lost (left) and gained (right) peaks overlapped with the predicted enhancer regions (from GeneHancer) and H3K27ac mark from relevant cancer cell lines (harbouring gain-of-function mutations in p53) for breast and colon cancers. (B) Heatmap shows the chromatin accessibility data centred at the gained peaks (and immediate flanking regions) that overlapped histone marks (including those that overlapped both histone marks and enhancer predictions) from MDA-MB-468 cell line, only the enhancer predictions or none in breast cancer. (C) Similar heatmap for the lost peaks in breast cancer.

Previous studies have shown that the wild-type and mutant p53 protein can recognize specific non-B-form DNA structures (23). To check this, we obtained the genome-wide prediction of different non-B-form structures and checked for their enrichment in the gained and lost peak regions (see Methods). This showed that only the G-quadruplex structure was significantly enriched in the gained and lost peaks, especially in the promoter regions, in breast carcinomas (and in gained peaks in colon adenocarcinoma) as compared to the random peaks in both tumour types (one-sided Fisher’s exact test, P < 0.001, see Supplementary Table S6).

### Impact of chromatin accessibility changes on gene expression

We then investigated if the differentially accessible peaks observed in the mutant p53 tumours lead to changes in the expression of the relevant genes. For this, we performed the differential RNA expression analysis between the wild-type and mutant p53 groups in breast and colon adenocarcinoma (see Methods) and compared the results with the genes annotated from the differential chromatin accessibility analysis. With the |log-fold change| > 1 and FDR (q-value) < 0.1, we identified 251 downregulated and 152 upregulated protein-coding genes in breast carcinoma (Figure 4A, Supplementary Table S7). Of these, six downregulated genes (EGR2, D4S234E, POU3F1, IL24, PENK, DUSP13) and three upregulated genes (ME1, KRT16 and PI3) were targets previously known to be regulated by mutant p53 protein (based on the literature) (24). In addition, 49 out of 251 downregulated and 19 out of 152 upregulated genes were also identified from the differential chromatin accessibility analysis (Figure 4C, Supplementary Table S8), including the known target genes IL24 and ME1. To further extend this, we focused on the genes identified from differential chromatin accessibility analysis that was supported by expression change in a similar direction (that is, upregulated -- gain in accessibility with expression log-fold change > 1, and downregulated -- a loss in accessibility with expression log-fold change < −1), and were not affected by CNA events (Figure 4C). Each gene was annotated with the presence of de novo motifs identified, p53 response element and other known co-factors of p53 binding to the differentially accessible regions of the respective genes (as shown in Figure 4C). Gene set enrichment analysis showed that some of the downregulated genes (such as GCM1, TP63, SDK1, NRG3, BCL2, NTRK2, EPHA7, TBX3, CNTN4, ROBO2, BMPR1B, VSTM2A) were enriched in cell differentiation and proliferation regulation. Of these, BMPR1B (for example) was found with a high number (n=53) of chromatin accessibility lost peaks across the gene body and enhancer regions (Figure 4E). BMPR1B encodes a member of bone morphogenetic proteins (BMPs) that belong to the transforming growth factor-β (TGF-β) family. It has been reported that reduced expression of BMPR1B can increase the proliferation of breast cancer cells, and is also associated with poor prognosis and bone metastasis in breast cancer patients (25, 26). Consistent with the loss of accessibility, we observed a lower expression of this gene in the mutant p53 samples as compared to the wild-type p53 samples (Wilcoxon rank-sum test, P = 0.08). We checked this association in additional TCGA breast cancer samples (which does not have chromatin accessible data) and found that the expression of BMPR1B was significantly lower in mutant p53 samples (Wilcoxon rank-sum test, P = 2×10^−14^) (Figure 4E). This suggests that in mutant p53 tumours the lower expression of BMPR1B can be explained by the changes in the chromatin structure, mediated by the mutant p53 protein. On the other hand, among the up-regulated genes, we found DAPK1 (Death-associated protein kinase 1) as one of the interesting candidates (Figure 4F). DAPK1 is a positive mediator of gamma-interferon induced programmed cell death. It has been shown that the high expression of DAPK1 can cause increased cell growth, and the depletion or inhibition of DAPK1 suppressed the cell growth, especially the mutant p53 but not wild-type p53 breast cancer cells (27). We observed an increase in accessibility in the enhancer regions of DAPK1 (Figure 4F), harbouring binding sites for the p53 response element, co-factors and G-quadruplex structure. The increase in DAPK1 expression was observed in these mutant p53 samples (P = 0.003), as well as in the additional TCGA breast cancer samples (P = 1×10^−8^). This suggests that the mutant p53 protein, together with other co-factors, could potentially transactive the expression of DAPK1 in these tumours.

**Figure 4:**
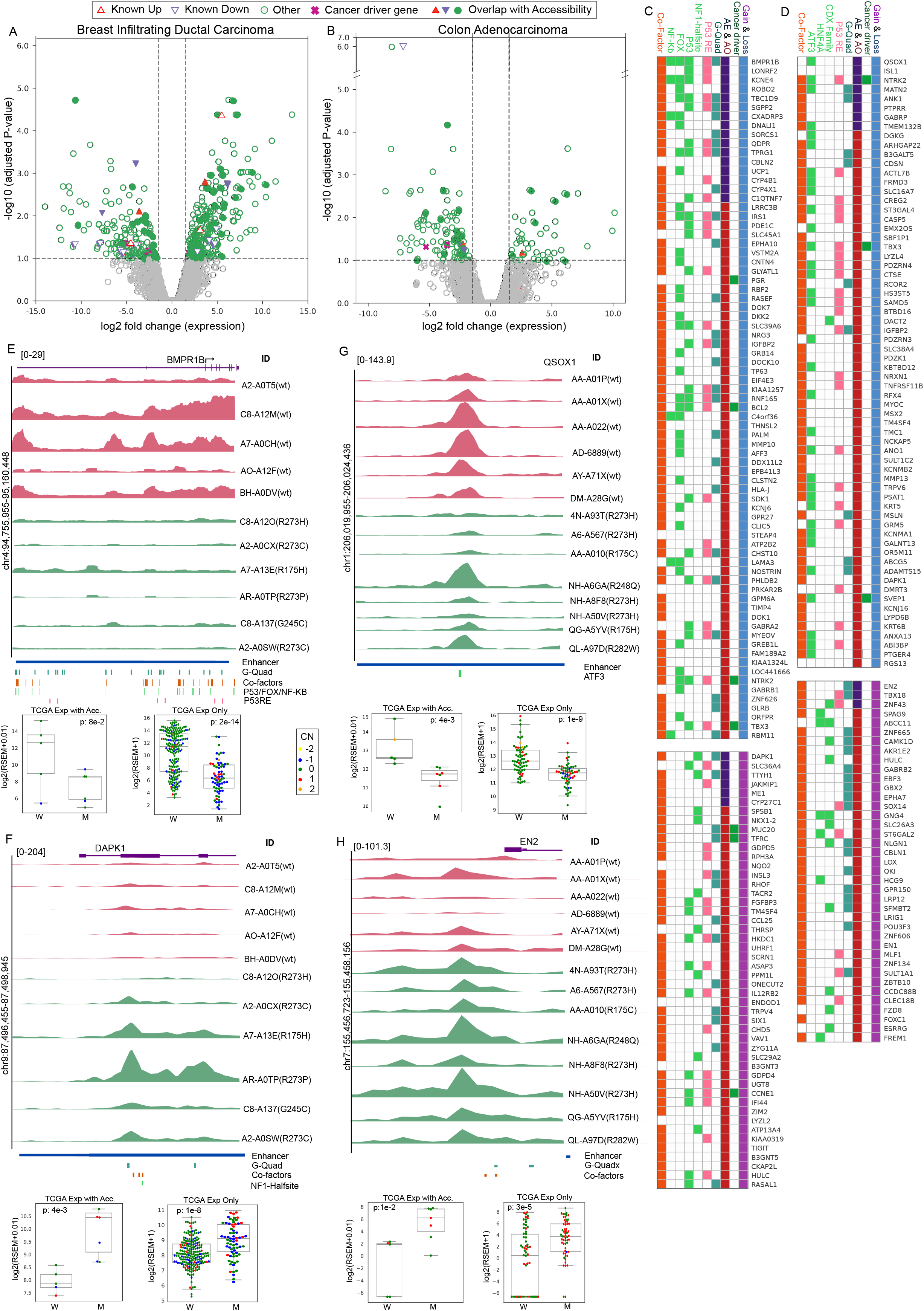
Impact of differential chromatin accessibility on gene expression. (A-B) Volcano plot shows the differentially expressed genes in mutant p53 with respect to the wild-type p53 tumours. Each dot represents a gene, and the x-axis represents the log2 fold change of expression value in mutant versus wild-type p53 tumours, and the y-axis shows the −log10 adjusted p-value. The vertical and horizontal dashed lines indicate the threshold of FDR (q-value) < 0.1 and |log2FoldChange(expression)| of 1. The triangle and inverted triangle symbol indicate the known target genes regulated by mutant p53, the symbol X indicates the cancer driver genes. The filled symbols indicate the genes that appeared significant in the differential accessibility analysis as well. (C-F) The chromatin accessibility profile in mutant and wild samples p53 samples for selected genes that showed significant differential gene expression and differential chromatin accessibility peaks. The plot was generated using SparK, each row represents a sample, and the annotation of predicted enhancer regions (from GeneHancer), predicted G-quadruplex motifs, and binding sites of known mutant p53 co-factors (See methods). The boxplot shows the expression of the gene in wild-type (W) versus mutant p53 (M) tumours for the matched samples with chromatin accessibility data and in additional TCGA samples which do not have chromatin accessibility data. Each dot represents a sample and the colour indicates the relative copy number status of the gene (−2 deep deletion, −1 deletion, 1 amplification, 2 high amplification) in the respective samples. The p-value was computed using the Wilcoxon rank-sum test. (G-H) The heatmap (generated using gitools) shows the list of genes that have differential accessible peaks and have expression change in the same direction (gain in accessibility with increased expression, and loss in accessibility with reduced expression). Each gene is annotated whether it overlapped with known cancer driver genes (from COSMIC), appeared in either differential expression analysis (AE), differential accessibility (AO) or both, presence of G-quadruplex, P53 response element, predicted de novo motifs and/or known mutant p53 co-factors in the differential accessibility region.

In colon adenocarcinomas, we found 95 downregulated and 44 upregulated genes from the differential RNA expression analysis. Of these, three downregulated genes (KIAA1324, ANK1, ABCA12) and one upregulated gene (ZNF415) were previously known targets of mutant p53 proteins (24). In addition, 13 out of 95 downregulated (including ANK1) and 3 out of 44 upregulated genes were also identified from the differential chromatin accessibility analysis (Figure 4D, Supplementary Table S8). The lesser number of overlap can be explained by the fact that two of the samples with chromatin accessibility data did not have matched gene expression data (see Methods). Further, we focused on the genes identified from differential chromatin accessibility analysis that showed expression change in a similar direction (as mentioned above) (Figure 4D). Gene set enrichment analysis showed that the upregulated genes, that has gain in the accessibility and increased expression (such as DMRT3, EBF3, EN1, EN2, ESRRG, FOXC1, GBX2, POU3F3, SFMBT2, SOX14, SPDEF, TBX18, TFAP2A, ZNF134, ZNF43, ZNF606, and ZNF665), were enriched in transcription regulatory activity and chromatin function. For example, the gene EN2 (Engrailed-2) is a member of the engrailed homeobox family. A recent study has shown the upregulation of EN2 in colorectal cancers associated with poor prognosis, and the knockdown of this in a colorectal cancer cell line, SW480 (harbouring GOF mutant p53), has inhibited proliferation and migration capacities of the cells (28). We observed a gained peak in the promoter region of this gene. This peak harbours a G-quadruplex structure and overlaps with the mutant p53 ChIP-seq derived from the SW480 cell line. Consistent with this, we observed a significant increase in expression of EN2 in mutant p53 samples in matched RNA seq data (P = 0.01) and in additional TCGA colorectal samples (P = 3 × 10^−5^) (Figure 4H). This suggests that the expression of EN2 can be regulated by the mutant p53 through direct binding, or together with their co-factors, in the promoter region. Among the downregulated genes, with loss of chromatin accessibility and reduced gene expression, we found QSOX1 as one of the interesting candidates. The loss of chromatin accessibility was observed at QSOX1 gene-enhancer interaction region located 28kb upstream of the TSS. This region also contains a predicted ATF3 motif (which was enriched in the lost peaks of colon adenocarcinoma, Figure 2G). QSOX1 helps in protein folding by oxidizing protein thiols by reduction of oxygen to hydrogen peroxide. It has been shown that downregulation of QSOX1 reduces cell-cell adhesion, and thus increases tumour migration and metastasis in breast cancer cell lines (29). QSOX1 expression has been shown to be a prognostic marker for breast and pancreatic cancer (30). However, a study on breast cancer cell-line (of epithelial origin) showed an inverse relationship of QSOX1 expression and cell proliferation (31). Consistent with this, we observed a significantly reduced expression of QSOX1 in matched samples (P = 0.004) and additional TCGA samples (P = 1 × 10^−9^) (Figure 4G).

In sum, by integrating the gene expression data, we identified some of the known and novel targets regulated by the mutant p53 potentially through chromatin changes. Some of these genes are involved in transcriptional regulation and regulation of cell differentiation and proliferation.

## Discussion

Several mechanisms have been proposed for the gain-of-function activity of mutant p53 under different cellular contexts and how that provides an advantage for the tumour cells to grow, metastasise and become resistant to treatments in different tumour types [see review (32)]. Although the previous studies including the ChIP-seq of mutant p53 and gene expression-based analysis helped to identify genes (dys)regulated in the mutant p53 cell lines and tumours, the local chromatin changes induced by mutant p53 (either through direct DNA binding or by interacting with other TFs or chromatin-modifying enzymes) in primary tumours is poorly understood. In this study, we tried to address this limitation by analysing the available chromatin accessibility data (ATAC-seq) generated in patient tumours from TCGA cohort, and identified differential accessible regions (that is, gained or lost chromatin accessible regions) in mutant p53 tumours compared to the wild-type p53 tumours, especially in breast infiltrating ductal carcinoma and colon adenocarcinoma. We found that the number of lost peaks are higher compared to the gained peaks in both cancer types, and this can be explained by the loss of wild-type p53 activity. However, the overlap of gained or lost peaks with the mutant p53 binding regions (inferred from ChIP-seq) data was less. This suggests that the mutant p53 preferentially binds to the existing open chromatin regions.

Moreover, we found that the gained and lost peaks were enriched for binding motifs for known co-factors of mutant p53 and other master TF regulators, enhancer marks and G-quadruplex structure (Figure 5). Taken together, this result suggests that the majority of the gained and lost peaks were located in the gene regulatory regions. By integrating gene expression data, we found that some of the genes linked to the differential accessible peaks showed significant expression change in matched tumour samples. This includes some of the known genes regulated by mutant p53 and other novel candidates. Compared to the previous studies that revealed mutant p53 specific gene expression signatures (based on the gene expression data alone) (33), this study provides additional information for those genes that underwent both local chromatin changes in the genic or regulatory regions, as well as the gene expression changes.

**Figure 5:**
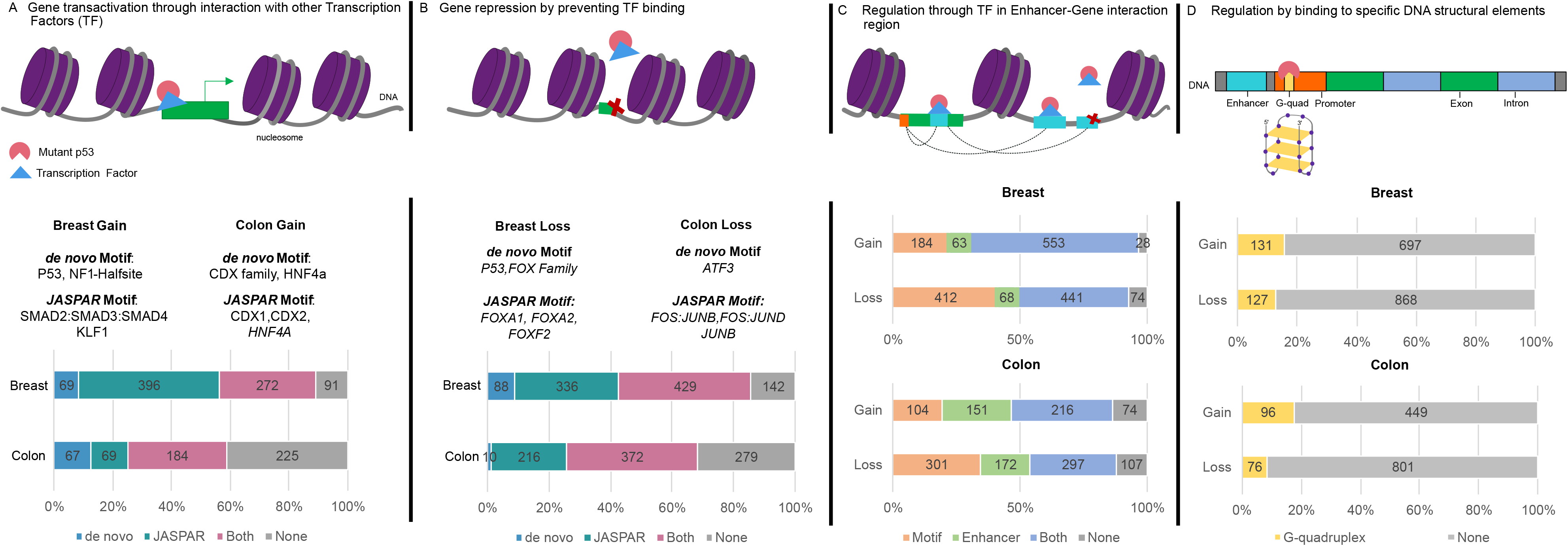
Model represents the possible regulation at differential chromatin accessibility regions. (A) The peaks that gained chromatin accessibility could be due to the transactivation by mutant p53 in combination with other TFs or interaction with chromatin-modifying enzymes, (B) The peaks that loss chromatin accessibility could be due to hijacking of TFs from the target regions by mutant p53 or interaction with chromatin-modifying enzymes, (C) enhancer-mediated gene regulation (which includes both (A) and (B)), and (D) direct gene regulation by binding of mutant p53 to specific DNA structural elements like G-quadruplex.

The limitations of the study include (a) although we integrated the chromatin accessibility, gene expression and somatic mutations data from the matched patient samples available in TCGA cohort, these data were not generated from the same cells and thus there could be some cell-to-cell variability, (b) we attempted to identify gained and lost peaks that potentially affected by the copy-number alteration. However, this can be further improved by incorporating allele-specific copy number data along with allele-specific chromatin accessibility data, and (c) given that many of the mutant p53 tumours analysed here have biallelic alteration in TP53 (that is, gain-of-function mutation together with loss of wild-type TP53 allelic), the differential chromatin accessibility peaks identified can be explained by the individual as well as combined effects of loss of wild-type p53 activity and GOF mutant p53 activity.

Overall, our analysis reveals tissue- and tumour-type specific chromatin changes in the mutant p53 tumours compared to the wild-type p53 tumours. Further studies are required to identify the mechanisms underlying the chromatin changes and the role of mutant p53 in it, and under different stress conditions.

## Materials and methods

### Identification of mutant p53 samples

We obtained the sample details of TCGA primary tumours (404 unique samples across 23 cancer types) that have chromatin accessibility data (ATAC-seq) from Corces & Granja et al. (13). For those samples, we retrieved the somatic mutation calls (MC3) (34) and somatic copy number data from the TCGA pan-can atlas study (https://gdc.cancer.gov/about-data/publications/pancanatlas). The allele-specific copy number estimated from the ASCAT algorithm (using Affymetrix SNP6 arrays data) was obtained from Martincorena et al (35). Further, we searched for samples harbouring gain-of-function (GOF) missense mutations in the DNA-binding domain of p53 (also referred as hotspot mutants), such as R273C/H/L/P/S/W/X/Y, R175A/C/G/H/L, V157F, Y220C, G245A/C/D/F/R/S/V, R248E/G/L/P/Q/W, R249G/K/M/S/W, R282G/H/Q/W and classified them as GOF mutant p53 samples (7). Samples that have both GOF missense mutation and minor allele copy number equals zero were classified as biallelic. Whereas, samples that do not carry any protein-affecting mutations (non-synonymous or nonsense) nor copy-number alterations in TP53 (that is, relative copy number value equals zero) were classified as wild-type p53 samples (see Table 1).

### Identification of differential chromatin accessibility regions

For the differential chromatin accessibility analysis, we focused on two specific cancer types (such as breast infiltrating ductal carcinoma and colon adenocarcinoma) that had sufficient sample size in the wild-type and mutant p53 groups (see Table 1). Cancer type-specific raw counts of ATAC-Seq peaks (called at the individual sample-level) were obtained from Corces & Granja et al. (13) (https://gdc.cancer.gov/about-data/publications/ATACseq-AWG). This matrix consisted of peaks on the rows and the sample information on the columns. From this, we selected the columns that matched the wild-type and mutant p53 samples. Further, to filter out peaks with low read counts, we converted the raw counts into CPM (counts per million) values to correct for the differences in library size between samples by using edgeR and removed peaks that do not have minimum 1 CPM in any of the samples considered (both wild-type and mutant). The remaining peaks (209,835 in breast infiltrating ductal carcinoma and 119,925 in colon adenocarcinoma) were subjected for differential analysis using the voom and limma eBayes packages. These tools were selected based on the recent benchmark study for the differential accessibility analysis using ATAC-seq data (36) and also used in Corces & Granja et al. (13). At first, the raw counts were transformed to log2(CPM) score (with a prior count of 5), followed by quantile normalization using voom. The resultant count matrix, along with weights obtained from the mean-variance relationship of each row (peak region), were fitted to a linear model using lmFit function. This model was then given to Limma eBayes function, along with the TP53 mutation status for each sample, to obtain the differentially accessible peaks in mutant p53 samples compared to the wild-type 53 samples. We performed this analysis separately for breast and colon cancers to avoid any bias from the tumour- and tissue-specific differences.

### Genomic annotation of peaks

The annotationPeaks function from HOMER (37) tool was used to annotate peaks to the closest gene possible (as well as obtained the gene-type information: protein-coding, ncRNA or pseudogene) and further categorize the location of the peaks as either promoter-TSS, 5’ UTR, exon, intron, 3’ UTR, TTS, non-coding or intergenic (based on the UCSC hg38 v6.4 annotations).

### Closest peak analysis

To find if the differentially accessible peaks are located close to the existing peaks, we computed the closest peak distance for each significant differentially accessible peak against non-significant peaks (from the respective cancer type) using bedtools closest function (with default parameters). The frequency of distance values was then binned into different groups <=1 kb, 1 to 2 kb, 2 to 5 kb, 5 to 10 kb, 10 to 15 kb, 15 to 30 kb, and > 30 kb (shown in Figure 1C and D).

### Copy number alteration annotation

The segmented copy number variation (SCNV) data from the Affymetrix SNP 6.0 array were downloaded from GDC portal (using TCGAbiolinks (38), especially the file type “grch38.seg.v2”). We intersected the genomic location of the differentially accessible peaks with the genomic location of segmented data (using bedtools (39)) and assigned the segment mean score to each peak. In cases where the peak overlapped with multiple segments, the median value of the overlapping segments was assigned. Then for each peak, we computed the median value of the segment mean score across wild-type and mutant p53 samples, respectively. Further, we annotated each peak that potentially has copy number gain if the median score > 0.3 in mutant or < −0.3 in wild-type, and copy number loss if median score < −0.3 in mutant or > 0.3 wild p53 samples. These annotations were shown in Figure 1E and F.

The gene-level relative copy number data was used to calculate a median score for mutant and wild sample groups (40). This was further used to filter out genes with copy number bias (median relative copy number was not equal to zero in both mutant and wild-type p53 samples) while selecting genes that showed both differential chromatin accessibility and expression change (Figure 4G, H).

### P53 response element and Mutant p53 ChIP data

To check whether any of the differentially accessible peaks contained a putative p53 response element, we used P53 retriever (with the default parameter) (19) to search for canonical (full site), as well as non-canonical (half-sites and 3/4 sites) p53 response elements.

Publicly available ChIP-seq data of mutant p53 from breast cancer cell lines HCC70 (harbouring GOF mutation R248) and MDA-MB-468 (R273)(16), and colon cell line SW480 (15)(R273), was obtained from ChIP-Atlas (41). We obtained the data from all these cell-lines and intersected with the differentially accessible peaks (using bedtools (39)) to annotate peaks that overlap mutant p53 binding sites (shown in Figure 1E and F). Further, to compare the peaks values between wild-type and mutant p53 samples, we computed the median CPM (across samples) for the mutant and wild-type p53 samples. The distribution of median CPM was shown in Figure 1G and H, and the difference between the wild-type and mutant median values for each peak (separated based on their genomic annotation) was shown in Supplementary Figure S1.

### Motif enrichment analysis

For TF motif enrichment analysis, we obtained the genome-wide transcription factor binding sites (TFBS) predictions based on curated TF motifs from JASPAR (with p-value < 0.0001) (42). The genomic coordinates of the gained and lost peaks were intersected with the TFBS to compute the percentage overlap (that is, the proportion of gained or lost peaks overlapping with the binding sites of a particular TF). Further, to check for the enrichment, we used one-sided Fisher’s exact test to compare the proportion of peaks overlapping with the binding sites of particular TF in gained or lost peaks with the proportion in randomly selected non-significant peaks (with the size and genomic annotations distribution equal to that of gained or lost peaks). The p-values were subjected to multiple hypothesis testing correction using Benjamini-Hochberg’s approach (Supplementary Table 3). Sites for previously known co-factors from JASPAR was also annotated in the significant peaks, these include: ELF1, ELF2, ELF4, GABPA, ERG, FLI1, FEV, ERF, ETV3, ELF3, ELF5, ESE3, ETS1, ETS2, SPDEF, ETV4, ETV5, ETV1, ETV2, SPI1, SPIB, SPIC, ELK1, ELK4, ELK3, ETV6, ETV7, SP1, SP5, SP2, AP1, HSF1, NFYA, NFYB, NFYC, TP63, TP73.

We also performed de novo motif enrichment analysis using HOMER’s findMotifsGenome (with default parameters) tool to search for sequence motif enriched in gained and lost peaks. To identify the motifs that are specifically enriched in the peaks that do not have predicted p53 response elements and/or overlap with the mutant p53 bindings, we split the gained and lost peaks based on the above criteria and performed the analysis separately in each group (for gain and lost peaks, Supplementary Figure S2). In both groups, we filtered out peaks that were annotated with copy-number alterations (gain or loss). The HOMER results for each group were shown in Table 5. The top k-mer motifs shown in Figures 2C and 2D were selected based on the enrichment p-value and percentage overlap.

The plotheatmap and plotprofile function from Deeptools (43) package was used to plot the accessibility profile centred at the peaks enrichment with particular TF motif or de novo motifs (shown in Figure 2).

### Enhancer and Histone enrichment

Curated Double Elite Gene-Enhancer interaction regions were obtained from GeneHancer database (44) and overlapped with the significant differentially accessible peaks in both cancer types. Additional information on active histone marks (such as H3K27ac) associated with enhancer regions was obtained from the respective cancer cells (harbouring GOF p53 mutations). SK-BR-3 (R175H), MDA-MB-468 (R273C), and AU565 (R175H) were selected for breast cancer; and SW480 (R273H), HT-29 (R273H), and NCI-N87 (R248Q) for colon cancer. The plotheatmap from Deeptools tools was then used to compare the accessibility profile of peaks that overlapped with both enhancers and histone marks, only enhancers or none.

### Non-B-form structure enrichment

A collection of predicted non-B DNA-forming motifs was obtained from non-B DB database (45). This includes A-Phased Repeats, Direct Repeat, G-Quadruplex Motif, Inverted Repeat, Mirror Repeat, Short Tandem Repeat, and Z DNA Motif. The genomic locations of these predicted non-B-form structure regions were intersected (using bedtools) with the gained or lost peaks to compute the percentage overlap and compared it with the randomly selected peaks (with size and genomic annotation ratio equal to that of gained or lost peaks) to compute the enrichment using one-sided Fisher’s exact test.

### Differential gene expression analysis

In breast, all samples with the chromatin accessibility data had matched gene expression data, but in colon two samples (one each from wild-type and mutant p53) didn’t have the matched expression data. The raw expression counts (rsem.genes.results) of protein-coding genes for breast and colon cancer samples were obtained from the GDC portal using TCGAbiolinks (38). Further, the count matrix was normalized (using TCGAanalyze_Normalization function) and filtered out genes with low expression count (using TCGAanalyze_Filtering with method = “quantile”, and qnt.cut = 0.25), leaving 14,892 genes in breast and 14,890 genes in colon. Further, the differential expression analysis was carried out using edgeR through TCGAanalyze_DEA function (with method = “glmLRT”). Genes with |log2foldchange| > 1 and FDR (q-value) < 0.1 were deemed significant.

The gene length normalized RSEM values was used to compute the median expression values for wild-type and mutant samples, and the expression fold change. The |log2foldchange| > 1 and neutral copy number in both mutant and wild p53 samples was used to filter genes from differential accessibility (as shown in Figure 4G and H). These genes were used for the gene set enrichment analysis using GSEA (46, 47). For the selected candidates in breast and colon, the expression difference in the matched TCGA samples, as well as in additional TCGA samples, was computed using the Wilcoxon rank-sum test.

## Acknowledgements

We thank Dimple Notani, Ramanathan Sowdhamini, S. Shivakumara, Núria López-Bigas, and members of RS lab for feedback and suggestions. We thank Eric Macwan for helping with the annotation of mutation status in patient samples. The results shown here are in part based upon data generated by the TCGA Research Network: https://www.cancer.gov/tcga. We acknowledge funding support from NCBS-TIFR and the Department of Atomic Energy, Government of India, under project no. 12-R&D-TFR-5.04-0800. RS acknowledges support from the Ramanujan fellowship (SERB, SB/S2/RJN-071/2018).

## Supplementary Figures and Tables

**Supplementary Figure S1**: The box-plot shows the distribution of the difference between the wild-type and mutant p53 accessibility scores (CPM) in peaks that overlapped the mutant p53 binding sites (inferred from ChIP-seq data). The peaks were split based on the genomics annotations shown on the x-axis.

**Supplementary Figure S2**: The significant gained and lost accessible peaks identified in breast and colon cancer have been categorized into those that overlapped regions with potentially copy number alteration, those that predicted to have p53 response element, overlap with mutant p53 binding sites (inferred from ChIP-seq data), both or none.

**Supplementary Figure S3**: The heatmap represents the overlap of differentially accessible peaks predicted with TF motifs from JASPAR and de novo motif analysis. The colour scale represents the Jaccard index score. The higher the value the more overlap between the peaks from different groups.

**Supplementary Table S1**: TCGA samples with TP53 mutation status.

**Supplementary Table S2**: The annotated significant differentially accessibility peaks identified in breast and colon cancers.

**Supplementary Table S3**: Enrichment of CNA events in different distance bins.

**Supplementary Tabel S4**: Results of TF motif enrichment analysis (related to Figure 2A-D).

**Supplementary Table S5**: Denovo motifs result from HOMER analysis of group A and B differentially accessible peaks.

**Supplementary Table S6**: Enrichment of non-B-form DNA structure in differentially accessible peaks.

**Supplementary Table S7**: Results from differential gene expression analysis.

**Supplementary Table S8**: Summary gene list that came up in the differential accessibility, differential expression, and overlap of both.

## Glossary

Mutant p53: TP53 gene with a hotspot (missense) mutation in the DNA-binding domain
Wild p53: TP53 gene with no protein-affecting mutations and copy number alterations
Lost peaks/region: Peak/region where chromatin accessibility has decreased in mutant p53 samples with respect to wild p53 samples
Gain peaks/region: Peak/region where chromatin accessibility has increased in mutant p53 samples with respect to wild p53 samples
GOF: Gain of Function
FDR: False discovery rate
CNA: Copy Number Alterations
p53 RE: predicted p53 Response Element
TF: Transcription Factor

